# Regulating oxygen avaibility mitigates oxidative stresses-induced antibiotic resistance gene expression in *E. coli* under climate warming

**DOI:** 10.1101/2025.06.06.658402

**Authors:** Wensi Zhang, Bharat Manna, Xueyang Zhou, Boyu Lyu, Naresh Singhal

## Abstract

Climate warming presents a critical challenge to global ecosystems, with rising temperatures inducing the emergence of antibiotic resistance genes (ARGs) in aquatic environments. Given the natural correlation between rising temperatures and declining oxygen levels in aquatic systems, we hypothesized that decoupling temperature and oxygen stress would reveal regulatory mechanisms mitigating ARG expression under future warming scenarios. Through transcriptomic analysis of Escherichia coli under temperature upshift (TU), dissolved oxygen upshift (DOU), and dissolved oxygen downshift (DOD), we identified 101 distinct regulons with 8 showing significant regulation (|log₂FC| > 1.5). Key regulons (*cusR*, *mhpR*, *pdhR*) exhibited parallel regulation under TU and DOD but opposite patterns under DOU, revealing mechanistic links between temperature and oxygen stress through shared regulatory networks involving oxidative stress regulators (*oxyR*-*metR*) and aerobic respiration control (*arcA*-*betI*). Evolved strains from controlled temperature-oxygen fluctuations demonstrated sophisticated metabolic reprogramming, including enhanced carbohydrate metabolism, nucleotide biosynthesis, and putrescine degradation. These strains exhibited optimized energy metabolism through reduced downregulation of ATP synthase subunits and NADH dehydrogenase complex genes, preserving electron transport chain function under thermal stress. Simultaneously, they displayed strategic ROS management with reduced upregulation of peroxiredoxin (*ahpF*) and Cu/Zn-SOD (*sodC*), while reversing Fe-SOD (*sodB*) expression patterns. To validate these mechanisms, we conducted independent batch reactor experiments with two dissolved oxygen conditions (2-8 mg/L) using intermittent perturbation patterns. Non-targeted metaproteomic analysis revealed that lowered oxygen levels (2 vs. 8 mg/L) triggered comprehensive reconfiguration of antimicrobial resistance mechanisms, including downregulation of penicillin-binding proteins, alanine racemase, and cationic antimicrobial peptide resistance systems, while modulating two-component regulatory networks. Our findings reveal coordinated regulatory networks optimizing energy metabolism, ROS management, and antibiotic resistance, suggesting oxygen management strategies could mitigate climate warming effects on antibiotic resistance in aquatic environments.

## Introduction

Climate warming is increasingly recognized as a pressing challenge, with profound extensive effects on terrestrial, marine, and freshwater ecosystems globally (Matthews & Weaver, 2010; Li et al., 2018). According to U.S. Environmental Protection Agency data, sea surface temperature has increased throughout the 20th century and continues to rise at an average rate of 0.14 °F per decade from 1901 to 2023 (Dunn et al., 2024). The end-of-century projections reveal catastrophic transformations in lake ice systems with +4.0 °C for lake temperature, −0.17 m for ice thickness and a 46-day decrease in ice duration relative to pre-industrial conditions under the most severe climate change scenario (Grant et al., 2021). Specifically, climate warming has been recognized as a major driver of bacterial resistance to antibiotics (Li et al., 2022). Antibiotic resistance genes (ARGs) are genetic elements that confer resistance to antibiotics in microorganisms, enabling them to survive and proliferate in the presence of antimicrobial agents (Wang et al., 2020). The widespread and prolonged misuse of antibiotics in clinical settings has led to the persistent presence of ARGs in various ecosystems (García et al., 2020). Climate warming may exacerbate the dissemination of ARGs through multiple mechanisms. Rising temperatures promote the proliferation of human pathogens, leading to increased infection rates and higher antibiotic consumption, which in turn exerts selective pressure favoring resistant strains (Wu et al., 2016). Additionally, elevated temperatures enhance horizontal gene transfer (HGT) among bacteria, a key process driving the spread of ARGs, thereby accelerating the evolution and dissemination of antibiotic resistance in microbial communities (Rzymski et al., 2024).

In response to these climate-driven resistance mechanisms, addressing the escalating crisis demands a comprehensive and multifaceted approach. From the perspective of source control, it is crucial to explore and identify novel therapeutic strategies or agents as alternatives to conventional antibiotics (Alaoui Mdarhri et al., 2022). Targeting specific cellular components, such as developing inhibitors for β-lactamases, efflux pumps, and outer membrane permeabilizers, also represents a promising strategy to combat antibiotic resistance (Martínez-Martínez, 2008). Moreover, effective disinfection remains a cornerstone strategy for limiting the dissemination of ARGs in aquatic environments (Zhu et al., 2023). Traditional water disinfection methods, such as chlorination, ultraviolet (UV) irradiation, the Fenton reaction, ozonation, and photocatalytic oxidation, have been widely applied to target antibiotic-resistant bacteria (ARB) and ARGs (Department et al., 2020). In recent years, innovative strategies have been developed to enhance the removal efficiency of ARGs, particularly in wastewater treatment plants (WWTPs). Among these, nanomaterial-based technologies have emerged as a promising biocontrol approach due to their exceptional adsorption capabilities (Kalia et al., 2020). However, these approaches face significant limitations, including high implementation costs, energy-intensive processes, and potential secondary environmental impacts. Their effectiveness requires further study and validation in complex natural water systems, where environmental variability and ecosystem interactions might hinder treatment performance. Consequently, there is an urgent need to explore nature-based strategies capable of mitigating the emergence of ARGs under climate warming conditions.

To develop these strategies, it is essential to understand how climate warming alters the metabolism of individual microbes in ways that contribute to antibiotic resistance. One emerging mechanism linking climate warming to antibiotic resistance involves oxidative stress responses in bacteria (X. Chen et al., 2019). Rising temperatures can lead to increased production of reactive oxygen species (ROS), causing oxidative stress and cellular damage in microorganisms (Sies & Jones, 2020). Strains that adapt to high temperatures show enhanced mechanisms for managing ROS, including upregulated antioxidant enzymes, improved DNA and proteins repair pathways (Seo et al., 2015). Research has begun to explore the role of oxidative stress response genes in antibiotic resistance, particularly under fluctuating environmental conditions. Studies have shown that oxidative stress response genes, such as *soxS*, *oxyR*, and *sodA*, play a critical role in mitigating the damage caused by reactive oxygen species (ROS) generated during antibiotic exposure (Dwyer et al., 2009). These genes help bacteria survive antibiotic-induced stress by repairing cellular damage and maintaining redox homeostasis (Ren et al., 2020). Experimental studies using untargeted metaproteomics have demonstrated increased expression of resistance proteins under elevated ROS levels in conventional activated sludge processes, suggesting a positive correlation between ARGs and reactive ROS (Manna et al., 2025). However, a significant gap persists regarding the molecular mechanisms underlying the relationship between climate warming and ARG expression and dissemination, specifically through oxidative stress response pathways and their regulatory networks in microbial metabolism.

Understanding these mechanisms requires examining how temperature and oxidative stress interact at the cellular level, as these environmental factors are intrinsically linked in aquatic systems. Rising temperatures typically lead to decreased oxygen concentrations in water (Matear & Hirst, 2003). Studies of *E. coli* transitioning between the outside environment and the mammalian GI tract found a highly significant overlap in sets of genes including TCA, cytochrome and heat shock genes downregulated by temperature upshift and oxygen downshift (Tagkopoulos et al., 2008). Given the natural correlation between rising temperatures and declining oxygen levels, we hypothesize that increasing oxygen availability may activate processes that are normally suppressed under heat stress which shares similarities with decreasing oxygen. This insight suggested that this approach could be used to mitigate ARGs in the natural environment under global warming. A study attempted to decouple the effects of temperature and oxygen shifts in *E. coli*, generating evolved strains through cyclical modulation of both variables. These evolved strains displayed enhanced growth compared to their parental counterparts, but the study did not explore the associated metabolic changes or the regulation of ROS and ARGs. This previous study cyclically modulated both temperature and oxygen to accelerate *E. coli* adaptation. However, changes in the temperature of waterbodies are not expected to occur rapidly. This raises the question of whether oxygen modulation alone could yield similar benefits. Furthermore, the oxygen modulations in these experiments, representing a transition from anoxic to oxic conditions, raised another critical question: How would increasing the frequency of these shifts affect *E. coli*’s adaptation?

To address these questions, we reanalyzed the dataset from Tagkopoulos et al. (2008), which includes transcriptome profiles of E. coli under three distinct environmental conditions: temperature upshift, dissolved oxygen downshift, and dissolved oxygen upshift. The natural correlation between temperature and dissolved oxygen was then decoupled by subjecting *E. coli* cultures to controlled fluctuations in both parameters simultaneously. Through this process, evolved strains were obtained and compared with their parental counterparts under elevated temperature stress. Additionally, we conducted a systematic experiment using a pure culture of *E. coli*, a model pathogen, to examine how oxygen modulation regulates ROS levels and suppresses ARG emergence. We expected this approach to reveal: (1) how oxidative stress response genes and ARGs are regulated under higher temperature stresses, and (2) the potential link between oxidative stress response and antibiotic resistance in *E. coli* under dynamic environmental conditions. This comprehensive strategy aimed to elucidate how oxidative stress response genes mediate the relationship between temperature-oxygen adaptation and antibiotic resistance in *E. coli* under future warming scenarios. Our findings can provide novel insights into mitigating antibiotic resistance in aquatic environments affected by climate change through oxygen management strategies.

## 2. Materials and methods

### 2.1. Transcriptome data collection

Transcriptomic datasets for *Escherichia coli* strain MG1655 under conditions of temperature upshift (TU), dissolved oxygen upshift (DOU), and dissolved oxygen downshift (DOD) were obtained from a previously published study (Tagkopoulos et al., 2008). *E. coli* cultures were subjected to controlled fluctuations in temperature and DO simultaneously to decouple the natural correlation between these parameters. This approach enabled the selection of evolved strains, which were subsequently compared with their parental counterparts to evaluate transcriptional responses under elevated temperature stress.

### 2.2. Batch reactor experiments with different DO perturbations

To investigate the potential role of oxygen perturbations in mitigating ROS and ARG stress in *E. coli*, batch reactor experiments were conducted using a pure strain of *E. coli*. Six DO conditions were applied: constant aeration at 2 mg/L (CA2) and 8 mg/L (CA8), continuous perturbation at 2 mg/L (CP2) and 8 mg/L (CP8), and intermittent perturbation at 2 mg/L (IP2) and 8 mg/L (IP8). CA2 and CA8 maintained steady DO levels at 1.8-2.2 mg/L and 7.8-8.2 mg/L, respectively. CP and IP both alternated between aerobic and anoxic phases, with DO fluctuating between 0-2.0 mg/L (CP2/IP2) and 0-8.0 mg/L (CP8/IP8) but differed in their perturbation patterns. Experiments were conducted in two identical acrylic cylindrical bioreactors (1 L working volume), operated in parallel at 20°C. Biological triplicates were included for each condition. The microbial reaction medium contained 3.84 g/L sodium bicarbonate as an inorganic carbon source and 1 mL of a trace element solution. Over a 48-hour period, DO levels were precisely controlled via an Arduino Mega2560-based microcontroller system (Banzi and Shiloh, 2022) and determined using a probe.

Non-targeted metaproteomic analysis was performed to assess microbial protein expression under experimental conditions. Three biological replicates per condition were collected, followed by protein extraction, quantification, and analysis using nanoscale liquid chromatography coupled to tandem mass spectrometry (nano LC-MS/MS). Additionally, intracellular metabolite profiling was conducted using gas chromatography-mass spectrometry (GC-MS) to evaluate metabolic shifts. Metabolite extraction was followed by methyl chloroformate derivatization, with both biological and technical replicates included to ensure analytical robustness. The metaproteomic and metabolomic analyses were conducted following the methodologies described in Zhou et al. (2025).

### 2.3. Bioinformatics analysis and data visualization

To visualize functional gene abundance across conditions, heatmaps were generated using z-score normalization in ComplexHeatmap package in R (v4.2.2). Pearson correlation analysis (p-value < 0.05) was conducted to evaluate associations between ROS genes and differentially abundant ARGs. Data visualization and statistical graphics were generated using the ggplot2 package in R (v4.2.2) and OmicStudio tools (https://www.omicstudio.cn/).

## 3. Results

### 3.1. Identification of regulons in *E. coli*

Understanding the regulatory mechanisms underlying bacterial stress responses is crucial for predicting how environmental changes will influence antibiotic resistance development in aquatic systems. To elucidate these complex regulatory networks and their role in mediating temperature-oxygen-ARGs interactions, we conducted a comprehensive analysis of transcriptional regulons in *E. coli* under temperature upshift (TU), dissolved oxygen upshift (DOU), and dissolved oxygen downshift (DOD). Transcriptome analysis of *E. coli* identified 101 distinct regulons, of which 8 showed upregulation or downregulation (|log₂FC| > 1.5) in response to TU, DOU, or DOD (Fig.1; TabS1). The *aidB* regulon plays a key role in DNA repair mechanisms (Landini et al., 1994), while the *cusR* regulon is involved in regulating electron transport processes (Yamamoto & Ishihama, 2005). The mhpR regulon, which controls the catabolism of aromatic compounds (Manso et al., 2011), exhibits significant changes in regulation under stress conditions. Additionally, the *metJ* regulon acts as a global repressor of methionine biosynthesis and other sulfur metabolism genes (Augustus & Spicer, 2011). The pdhR regulon modulates pyruvate dehydrogenase complex activity, linking central carbon metabolism with energy production and responding to metabolic fluctuations (Göhler et al., 2011). Notably, we identified important regulatory network connections through two key regulons. *metR*, which is under the control of the oxidative stress regulator *oxyR* (Bogard et al., 2012), and *betI*, regulated by the aerobic respiration control protein *arcA*, highlighted the interconnected nature of temperature and oxidative stress responses (Lamark et al., 1996). Analysis of regulon regulation under three conditions revealed coordinated responses to temperature and oxygen stress. Notably, several regulons displayed similar regulation trends under TU and DOD conditions but showed opposite regulation under DOU. Specifically, *cusR* was downregulated in both TU and DOD conditions but upregulated during DOU. Similarly, *mhpR* and *pdhR* showed upregulation in TU and DOD conditions but downregulation in DOU. These parallel regulation patterns between temperature stress and low oxygen conditions suggest a mechanistic link between these environmental stressors. This coordinated regulation may represent an adaptive strategy where cells can leverage oxygen-responsive pathways to enhance their tolerance to high-temperature stress, particularly through the modulation of metabolic processes. The identification of these regulatory networks provides crucial insights into how *E. coli* integrates different stress signals to coordinate an effective response to temperature and oxygen stresses.

**Figure 1.**
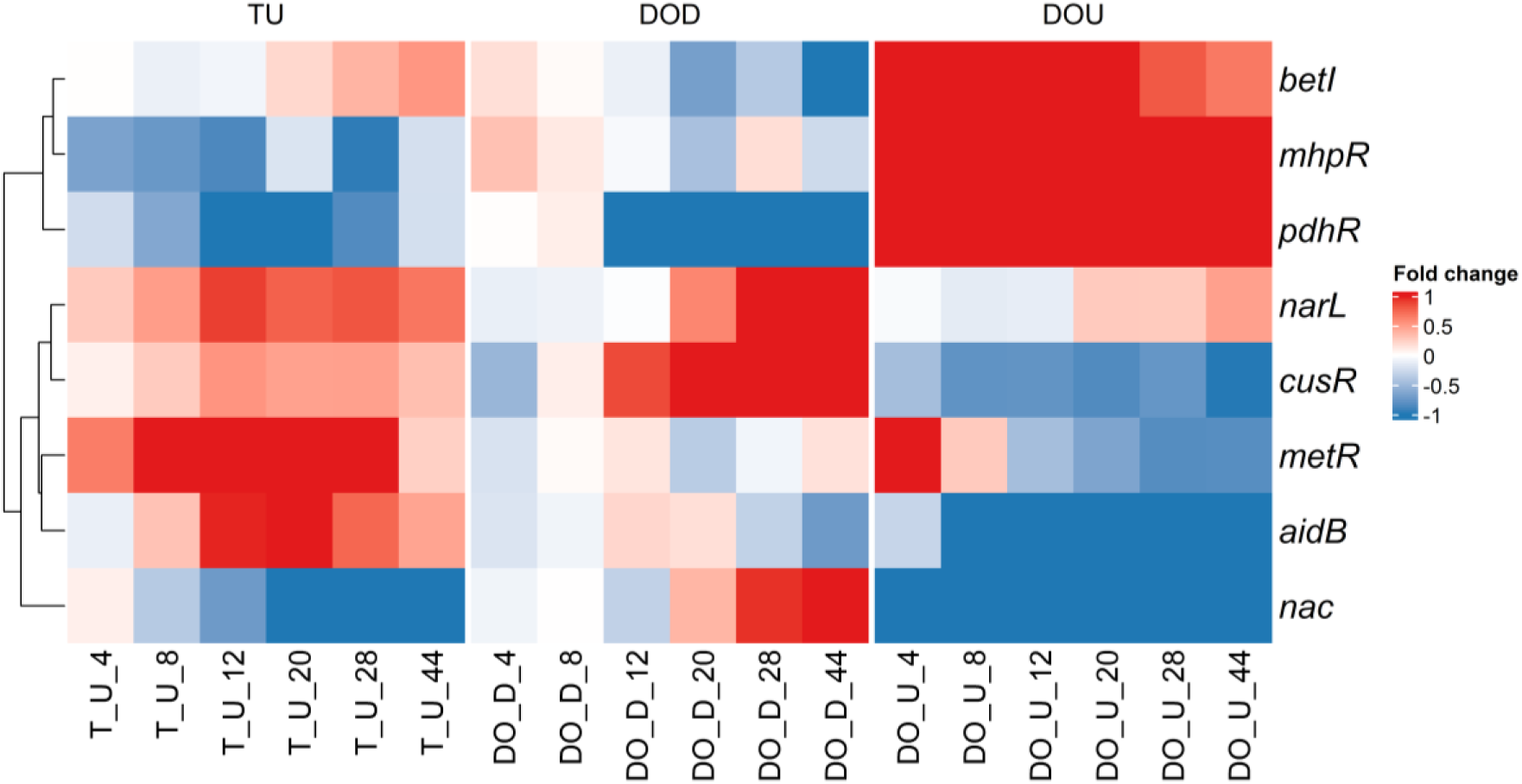
The fold change of eight regulons under temperature and oxygen stress conditions. Regulons were analyzed under temperature upshift (TU), dissolved oxygen upshift (DOU), and dissolved oxygen downshift (DOD) treatments, with numbers indicating time points treatment. Eight regulons showed significant regulation with |log₂FC| > 1.5 threshold.

### 3.2. Metabolic adaptations between evolved and parental *E. coli* strains

The regulatory network analysis revealed the foundational stress response mechanisms present in *E. coli*, but the question remained whether prolonged exposure to combined temperature-oxygen stress would lead to evolutionary adaptations that enhance these responses. To investigate this evolutionary dimension, we compared the transcriptome profiles of evolved strains with their parental counterparts under elevated temperature stress, focusing on how the regulatory networks identified might be modified or enhanced through adaptive evolution. The evolved strains exhibited significant upregulation of key metabolic pathways compared to parental strains, specifically in carbohydrate metabolism (respiration), amino acid degradation (particularly putrescine degradation), and nucleotide biosynthesis (Fig.2). These metabolic changes align with and extend beyond the regulatory responses identified in our regulon analysis. Enhanced respiratory metabolism could support increased energy demands for higher temperature stress response mechanisms (Kaila & Wikström, 2021). Meanwhile, upregulated nucleotide biosynthesis may facilitate DNA repair processes, complementing the DNA repair mechanisms controlled by the *aidB* regulon identified in our regulatory network analysis (Wozniak & Simmons, 2022). The enhanced putrescine degradation pathway is particularly noteworthy, as polyamine catabolism has been established as a crucial metabolic response to multiple stressors (Nair et al., 2025). Previous studies have demonstrated that strains lacking catabolic polyamine pathways show impaired responses to oxidative stress, high temperature, and antibiotic exposure, highlighting the fundamental role of polyamine metabolism in stress adaptation (Schneider et al., 2013).

**Figure 2.**
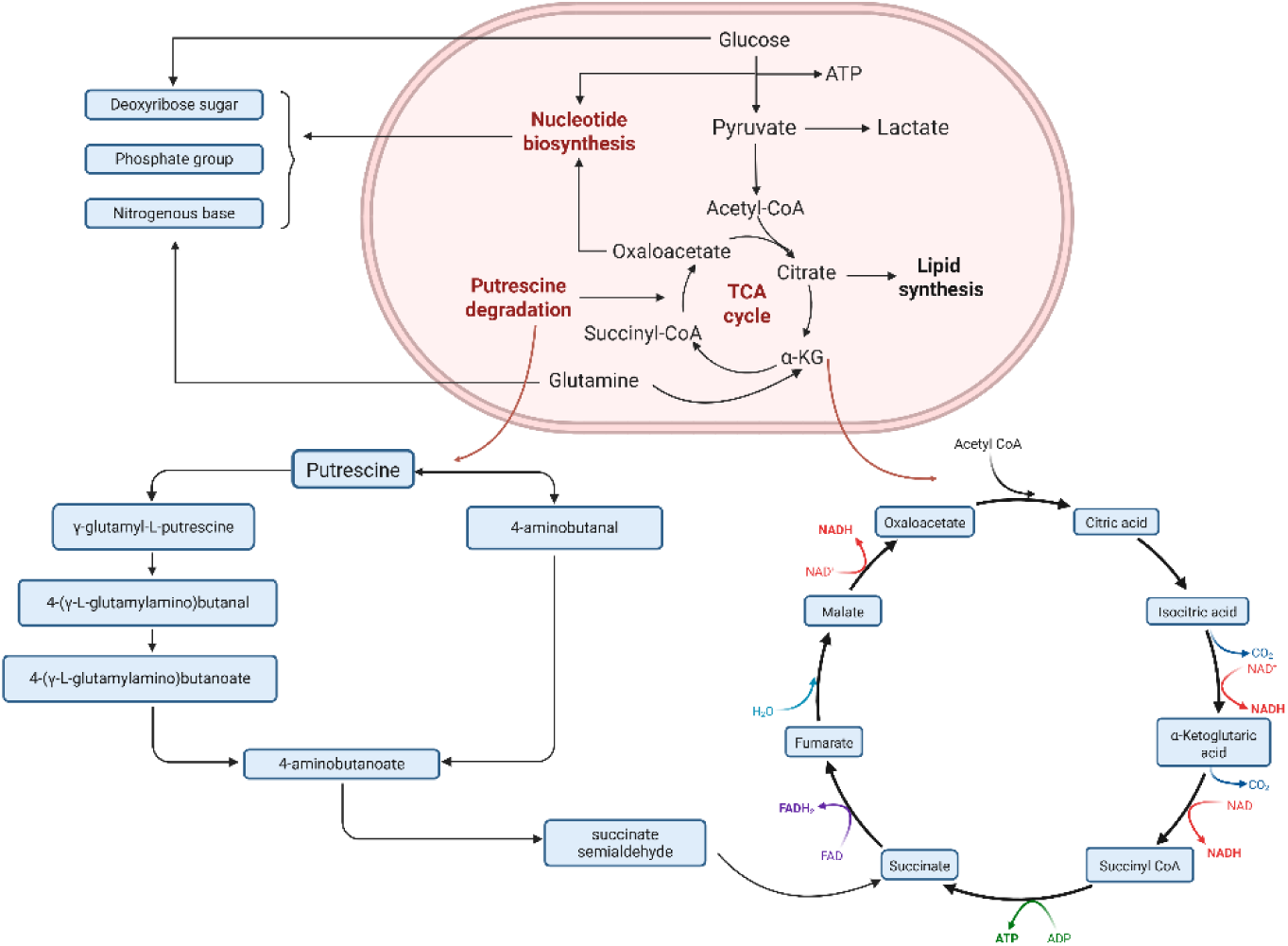
Comparison of globally regulatory genes (log₂FC > 1.5) between evolved and parental strains.

### 3.3 Comparison of ROS relevant between evolved and parental *E. coli* strains

#### 3.3.1. ROS scavenging genes

Transcriptome analysis revealed distinctive adaptations in oxidative stress management systems between parental and evolved *E. coli* strains under higher temperature stresses (Fig.3). The primary transcription regulator *oxyR* showed nearly identical average responses between parental and evolved strains. In contrast, *sodA* exhibited significantly less downregulation in evolved strains (log₂FC: −0.391) compared to parental strains (log₂FC: −1.240), suggesting enhanced maintenance of Mn-SOD activity at elevated temperatures (Touati, 1983). Conversely, genes typically upregulated during oxidative stress showed markedly reduced responses in evolved strains. The peroxiredoxin gene *ahpF* exhibited substantially less upregulation in evolved strains (log₂FC: 0.038) compared to parental strains (log₂FC: 0.403). Similarly, *sodC* showed reduced upregulation in evolved strains (log₂FC: 0.103) compared to parental strains (log₂FC: 0.360). Most notably, *sodB* displayed a complete reversal in expression pattern, shifting from downregulation in parental strains (log₂FC: −0.220) to upregulation in evolved strains (log₂FC: 0.134). The overall pattern of reduced oxidative stress gene regulation in evolved strains compared to parental strains suggested that evolved bacteria may experience lower baseline ROS stress levels, potentially due to enhanced metabolic efficiency or improved cellular antioxidant capacity developed through adaptive evolution.

**Figure 3.**
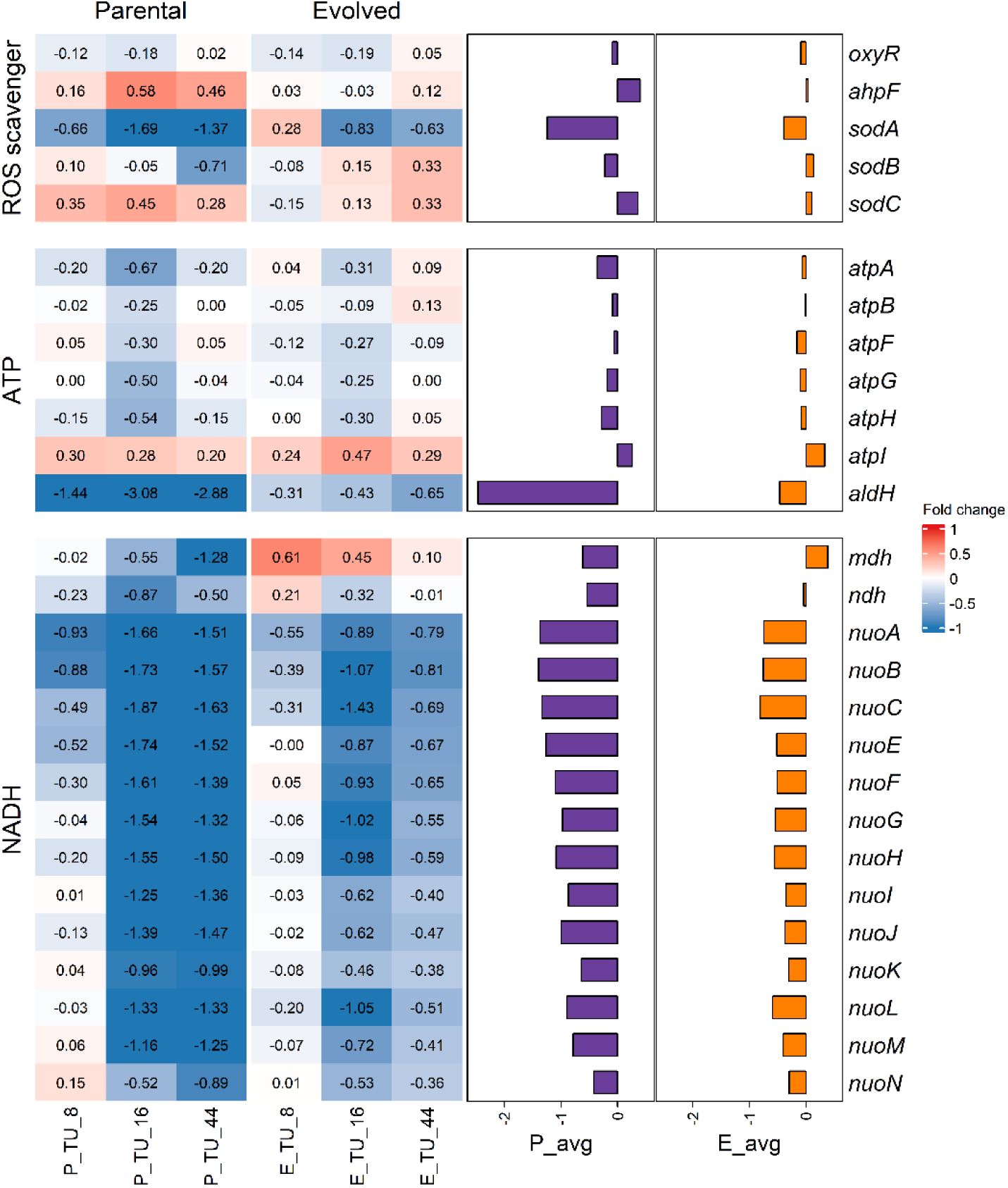
Fold change of ROS-related genes in parental and evolved strains under temperature upshift conditions. Heatmap shows fold change s between parental (P) and evolved (E) strains under temperature upshift (TU) treatment, with numbers indicating time points treatment. Adjacent bar chart displays average fold change values for each gene across all conditions, where P_avg represents the parental strain average and E_avg represents the evolved strain average.

#### 3.3.2. The ATP-related genes between the parental and evolved strains

Comparison of ATP synthase subunit genes showed that evolved strains consistently exhibited reduced downregulation compared to parental strains across most subunits (Fig.3). The most notable differences were observed in *atpA* (parental: log₂FC: −0.355 vs. evolved: log₂FC: −0.061), *atpH* (parental: log₂FC: −0.281 vs. evolved: log₂FC: −0.080), and *atpG* (parental: log₂FC: −0.181 vs. evolved: log₂FC: −0.097). The *atpB* gene showed minimal downregulation in both strains (parental: log₂FC: −0.087 vs. evolved: log₂FC: −0.005). Notably, *atpF* was the only gene showing greater downregulation in evolved strains compared to parental strains (parental: log₂FC: −0.062 vs. evolved: log₂FC: −0.158). Most remarkably, *atpI* demonstrated upregulation in both strain types but was more strongly upregulated in evolved strains (parental: log₂FC: 0.258 vs. evolved: log₂FC: 0.334).The reduced suppression of core ATP synthase subunits (*atpA*, *atpB*, *atpG*, *atpH*) in evolved strains indicates maintained or improved energy generation capacity during temperature stress, which aligns with the enhanced respiratory metabolism observed in our previous metabolic pathway analysis. This preservation of ATP synthase function might support the increased energy demands associated with stress response mechanisms and cellular maintenance at elevated temperatures, representing a key metabolic adaptation that enables evolved strains to better cope with thermal stress conditions.

#### 3.3.3. The NADH-related genes between the parental and evolved strains

Comparison of the NADH dehydrogenase I complex (encoded by nuo genes) also showed that evolved strains consistently exhibited less severe downregulation than parental strains across all subunits (Fig.3). The most pronounced differences were observed in *nuoB* (parental: log₂FC: −1.395 vs. evolved: log₂FC: −0.756), *nuoC* (parental: log₂FC: −1.333 vs. evolved: log₂FC: −0.809), and *nuoE* (parental: log₂FC: −1.263 vs. evolved: log₂FC: −0.514). Other *nuo* genes demonstrated similar patterns of reduced repression in evolved strains, with differences ranging from *nuoN* (parental: log₂FC: −0.420 vs. evolved: log₂FC: −0.294) to *nuoF* (parental: log₂FC: −1.101 vs. evolved: log₂FC: −0.508). The alternative NADH dehydrogenase ndh showed a dramatic improvement in evolved strains (parental: log₂FC: −0.533 vs. evolved: log₂FC: −0.043). Among related enzymes, aldehyde dehydrogenase (*aldH*) exhibited a striking contrast between strain types, with evolved strains showing substantially less repression (parental: log₂FC: −2.464 vs. evolved: log₂FC: −0.463). Most remarkably, malate dehydrogenase (*mdh*) displayed a complete reversal in expression pattern between strains, shifting from downregulation in parental strains (log₂FC: −0.614) to upregulation in evolved strains (log₂FC: 0.385). The consistent pattern of reduced downregulation in NADH dehydrogenase complex genes indicates that evolved strains have developed enhanced capacity for maintaining electron transport chain function under thermal stress compared to their parental counterparts. This differential regulation pattern suggested that evolved strains have developed superior mechanisms for preserving energy metabolism and redox balance during temperature stress. The coordinated maintenance of these critical respiratory components in evolved strains supported enhanced ATP production capacity, forming an integrated adaptation for improved metabolic resilience under thermal stress conditions.

### 3.4. Comparison of ARG between evolved and parental *E. coli* strains

Transcriptome analysis revealed significant adaptations in antibiotic resistance gene (ARG) expression between parental and evolved *E. coli* strains under temperature stress (Fig.4). The evolved strains exhibited substantial reduction in expression of the multiple antibiotic resistance regulator *marA* (log₂FC: 0.099 vs 0.481 in parental strains), indicating a strategic reduction in broad-spectrum antibiotic resistance mechanisms. Similar trends were observed across membrane modification genes (particularly the *arn* family), which showed an average reduction of 0.227 log₂FC: in evolved strains, suggesting decreased lipopolysaccharide modifications associated with antibiotic resistance. The alanine racemase gene (*alr*), essential for peptidoglycan synthesis and β-lactam resistance, displayed significantly less downregulation in evolved strains (log₂FC: −0.065) compared to parental strains (log₂FC: −0.447). Similarly, the ampG permease gene showed remarkable adaptation (log₂FC: −0.020 vs −0.514), suggesting maintained cell wall integrity without sustained antibiotic resistance upregulation. The two-component systems involved in stress response and antibiotic resistance (*phoP/Q* and *basS*) also showed modulated expression in evolved strains. Notably, phoP displayed downregulation in evolved strains (log₂FC: −0.265) compared to slight upregulation in parental strains (log₂FC: 0.029).

**Figure 4.**
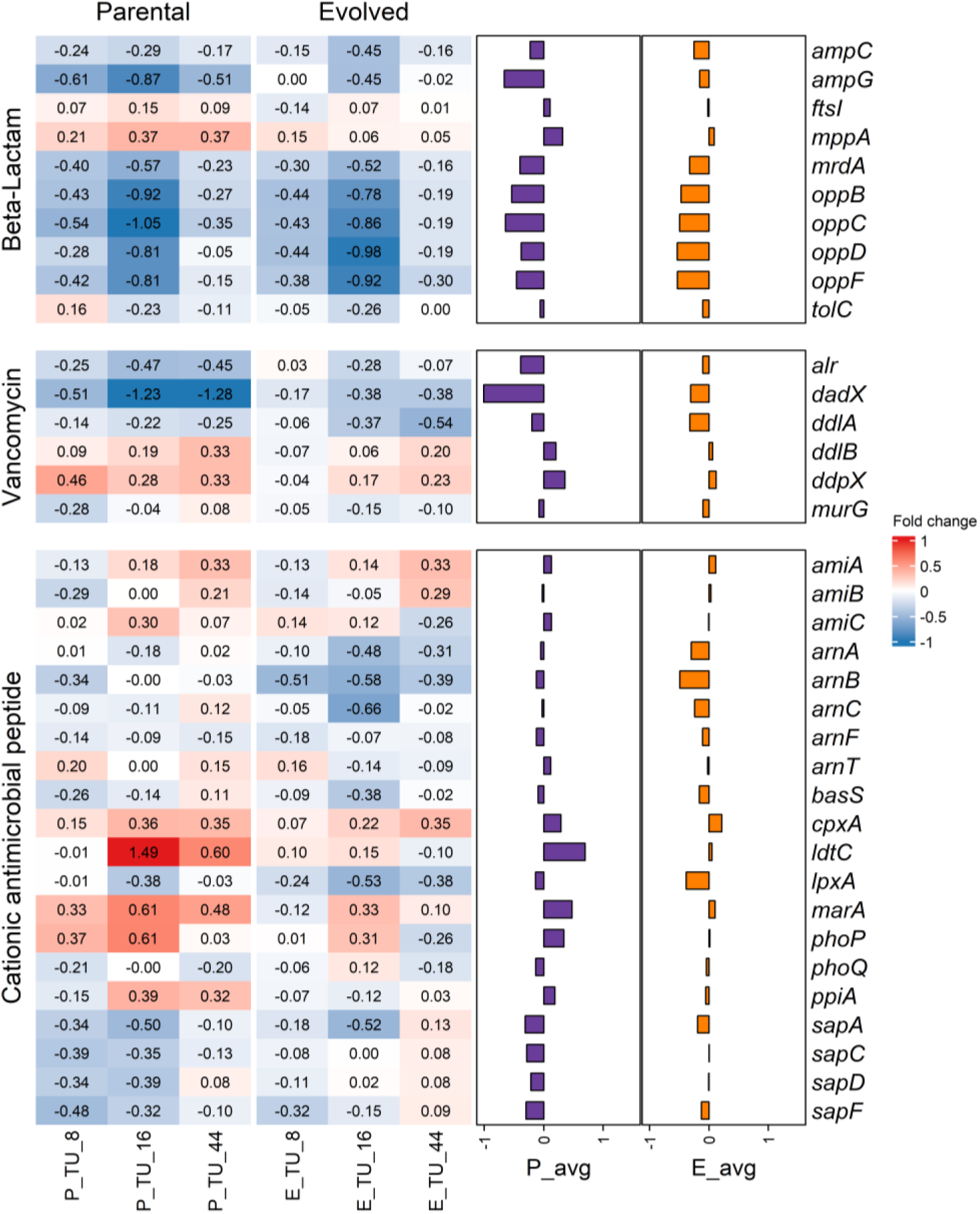
Fold change of ARGs in parental and evolved strains under temperature upshift conditions. Heatmap shows fold change s between parental (P) and evolved (E) strains under temperature upshift (TU) treatment, with numbers indicating time points treatment. Adjacent bar chart displays average fold change values for each gene across all conditions, where P_avg represents the parental strain average and E_avg represents the evolved strain average.

These transcriptomic adaptations collectively suggest that evolved strains have developed more efficient stress response mechanisms that reduce reliance on energy-intensive antibiotic resistance pathways while maintaining cellular integrity under thermal stress. The evolved strains demonstrated enhanced antibiotic tolerance, characterized by reduced regulation of ARGs compared to the parental strains, indicative of a more efficient and streamlined stress response system. This improved tolerance aligns with key resistance mechanisms observed across various antibiotics. These findings highlight the interplay between evolved resistance mechanisms and stress response efficiency in bacterial survival and adaptation.

### 3.5. Validation of ROS scavenger genes under low and high oxygen perturbation conditions

Proteomic analysis demonstrated differential expression of oxidative stress response proteins between low (2 mg/L, IP2) and high (8 mg/L, IP8) oxygen perturbation conditions (Fig.5). The antioxidant enzymes alkyl hydroperoxide reductase subunits C and F (*AhpC* and *AhpF*) showed consistent downregulation under high oxygen conditions, with average log₂FCs of −0.970 and −0.910, respectively. Similarly, superoxide dismutase (*SodA*) exhibited the strongest downregulation among all examined proteins with an average log₂FC of −1.360 in high oxygen conditions. Catalase proteins displayed modest changes in abundance, with both *KatE* and *KatG* showing slight average downregulation (−0.200 and −0.340 log₂FC, respectively) under increased oxygen exposure. Notably, the transcriptional regulator OxyR demonstrated upregulation with an average log₂FC of 0.680, although with substantial variability across replicates.

**Figure 5.**
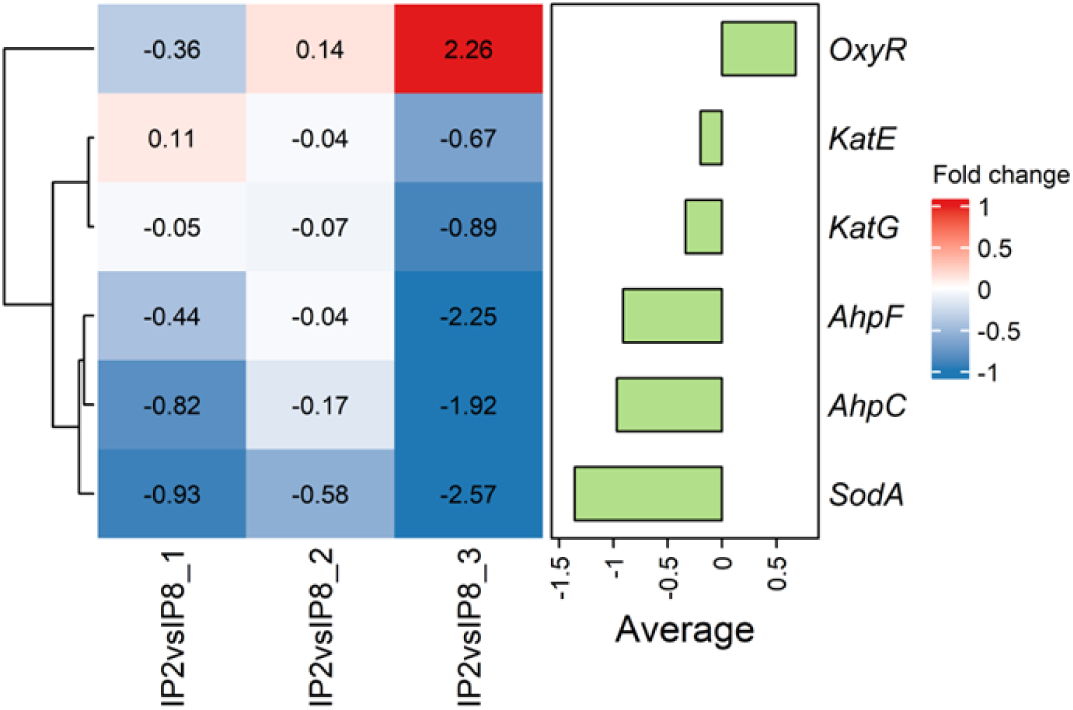
Differential expression of ROS scavenger genes between low (2 mg/L, IP2) and high (8 mg/L, IP8) oxygen perturbation conditions (Log₂FC) in *E. coli*. Adjacent bar chart displays average fold change values for each gene across all conditions.

The proteomic profile of oxidative stress response proteins indicates a paradoxical adaptive response to elevated oxygen levels. Surprisingly, the downregulation of *AhpC* and *AhpF* under high oxygen conditions contradicts the expected induction of antioxidant defenses. This reduction in peroxide-detoxifying enzymes, along with the strong decrease in *SodA* abundance, suggested a potential acclimation mechanism where cells may be adjusting their redox homeostasis systems to the consistently higher oxygen environment. The modest downregulation of both catalase proteins (*KatE* and *KatG*) further supports this adaptive response pattern. Interestingly, the transcriptional regulator *OxyR* showed overall upregulation despite the downregulation of its target proteins. This apparent contradiction may represent a complex regulatory mechanism where increased *OxyR* serves to maintain basal expression of antioxidant enzymes while preventing their overexpression, which could be metabolically costly under sustained high oxygen conditions. Collectively, these proteomic results suggest that exposure to higher oxygen levels (8 mg/L vs. 2 mg/L) leads to metabolic adaptation rather than a typical stress response. This may reflect the organism’s ability to acclimate to different oxygen regimes by recalibrating its antioxidant defense systems to optimize energy expenditure while maintaining effective protection against oxidative damage.

### 3.6. Validation of ARGs under low and high oxygen perturbation conditions

Proteomic analysis was conducted to examine the differential expression of antimicrobial resistance genes between low (2 mg/L, IP2) and high (8 mg/L, IP8) oxygen perturbation conditions. The data revealed distinct expression patterns across three major antimicrobial resistance pathways: beta-lactam resistance, vancomycin resistance, and cationic antimicrobial peptide (CAMP) resistance.

Beta-lactam resistance proteins showed predominantly negative fold changes, indicating downregulation in high oxygen conditions. Cell division protein FtsI (penicillin-binding protein 3) showed consistent downregulation across replicates (average log₂FC: −1.90). Penicillin-binding proteins 1A and 2 (*MrcA* and *MrdA*) also exhibited substantial downregulation in at least one replicate. The oligopeptide transport system components (*OppA*, *OppD*, *OppF*) showed variable expression patterns, with *OppF* consistently downregulated (average log₂FC: −1.09). Among vancomycin resistance proteins, alanine racemase (*Alr*) showed strong downregulation (average log₂FC: −2.06), while its isoform DadX showed modest downregulation only in one replicate. D-alanine-D-alanine ligases (*DdlA* and *DdlB*) and UDP-N-acetylmuramoyl-tripeptide--D-alanyl-D-alanine ligase (*MurF*) showed minimal changes in expression. UDP-N-acetylglucosamine transferase (*MurG*) exhibited moderate downregulation (average log₂FC: −0.38). The CAMP resistance pathway showed the most variable expression patterns. Notably, signal transduction protein *PmrD* was consistently downregulated across all replicates (average log₂FC: −2.13). KDO II ethanolaminephosphotransferase (*EptB*) showed extreme variability, with strong upregulation in the first two replicates but strong downregulation in the third. The two-component system sensor histidine kinase *PhoQ* was upregulated in two replicates (average log₂FC: 1.51), while its response regulator *PhoP* showed modest upregulation across all replicates (average log₂FC: 0.27). The cationic peptide transport system components showed mixed patterns, with *SapC* consistently and strongly downregulated (average log₂FC: −2.78).

The proteomic analysis revealed a complex regulatory network whereby elevated oxygen levels orchestrate a comprehensive reconfiguration of antimicrobial resistance mechanisms in E. coli. Under high oxygen conditions, the observed downregulation of key cell wall synthesis machinery, including penicillin-binding proteins (*FtsI*, *MrcA*, *MrdA*) and peptidoglycan synthesis enzymes, likely represents an energy conservation strategy that reduces metabolically expensive cell wall remodeling processes. Concurrently, the variable expression patterns of lipopolysaccharide modification enzymes (*ArnA*, *EptA*, *EptB*) indicate adaptive restructuring of outer membrane architecture, potentially modulating resistance to cationic antimicrobial peptides through altered membrane charge distribution. This adaptive response is further coordinated through a reconfiguration of two-component regulatory systems, where the upregulation of *PhoQ*/*PhoP* paired with *PmrD* downregulation suggested precise fine-tuning of resistance gene expression to match oxygen-specific stress conditions. Additionally, the consistent downregulation of transport system components (*SapC*, *OppF*) indicates reduced peptide uptake and efflux capacity, which may simultaneously limit nutrient acquisition while modifying antimicrobial resistance profiles. These proteomic changes demonstrate that oxygen availability serves as a master regulator that coordinates multiple cellular systems to optimize survival under aerobic stress conditions while potentially reducing the metabolic burden associated with maintaining high-level antimicrobial resistance.

These findings suggest that increased oxygen levels trigger a reprogramming of antimicrobial resistance mechanisms, potentially reflecting a trade-off between energy conservation and defense capability. The observed downregulation of key resistance proteins may indicate reduced antibiotic tolerance under high oxygen conditions, which could have important implications for antibiotic treatment strategies. Alternatively, these changes may represent a shift from constitutive to inducible resistance mechanisms, where resources are conserved until specific antimicrobial threats are detected. The variability observed across replicates highlights the complexity and potential stochasticity of these adaptive responses.

**Figure 6.**
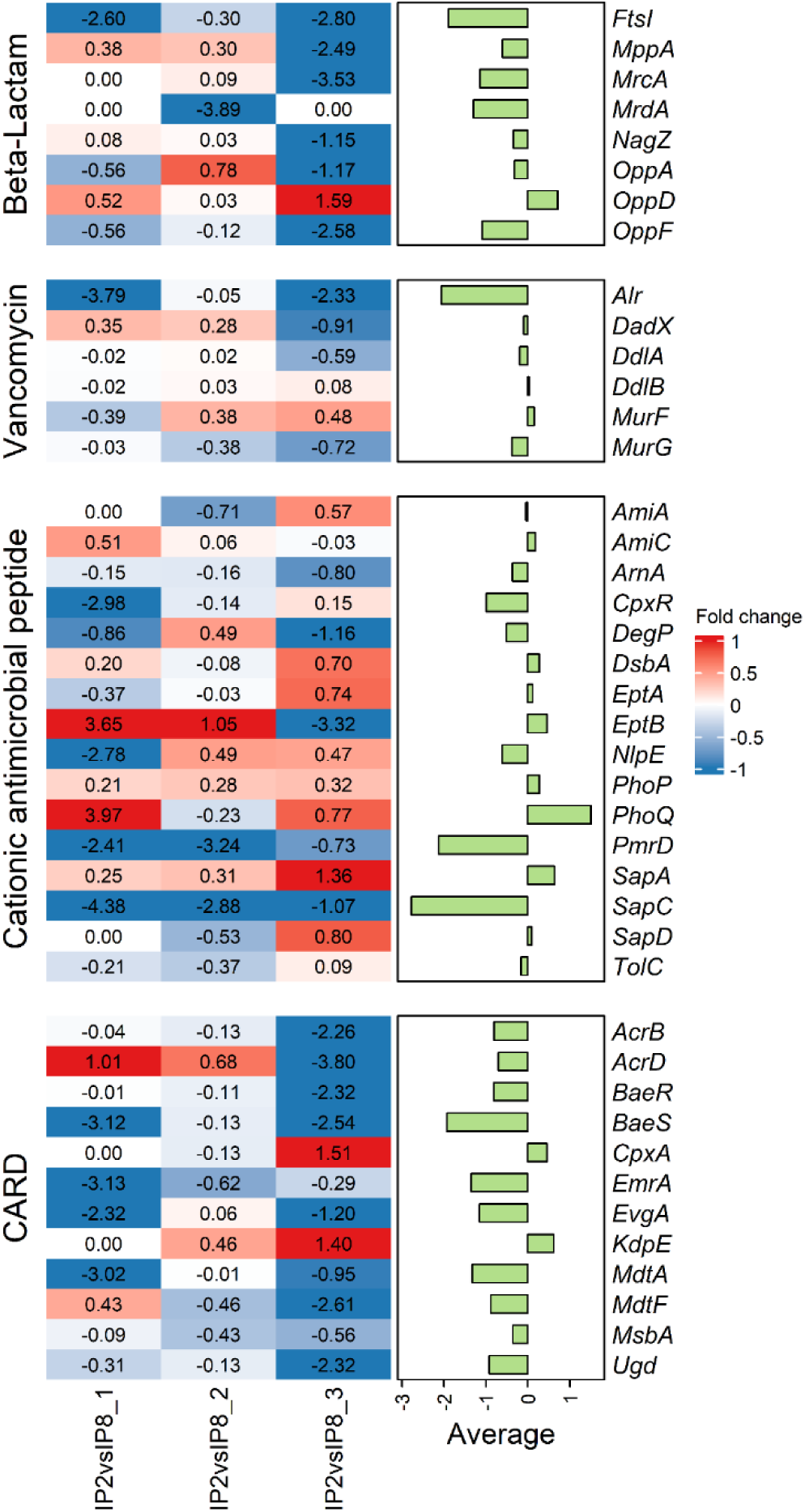
Differential expression of ARGs between low (2 mg/L, IP2) and high (8 mg/L, IP8) oxygen perturbation conditions (Log₂FC) in *E. coli*. Adjacent bar chart displays average fold change values for each gene across all conditions.

## 4. Discussion

### 4.1. Integrated regulatory networks drive metabolic adaptation under higher temperature stress

Our findings reveal a sophisticated regulatory framework that enables *E. coli* to coordinate responses to higher temperatures stressors through interconnected transcriptional networks. The identification of 8 regulons with coordinated responses to temperature and oxygen stress provides crucial insights into bacterial adaptation strategies under climate warming scenarios. The parallel regulation patterns observed in key regulons (*cusR, mhpR, pdhR*) under temperature upshift and dissolved oxygen downshift, contrasted with their opposite regulation under dissolved oxygen upshift, demonstrate that bacteria can leverage existing oxygen-sensing pathways to enhance thermal stress tolerance (Seo et al., 2015). This regulatory convergence suggests an evolutionary optimization where environmental stress responses are integrated through shared molecular mechanisms, particularly through oxidative stress regulators (*oxyR-metR*) and aerobic respiration control (*arcA-betI*) networks (Chen et al., 2018; López-Maury et al., 2008).

The metabolic reprogramming observed in evolved strains represents a fundamental shift from reactive damage repair to proactive stress mitigation, addressing the research gap identified regarding molecular mechanisms underlying climate warming and antibiotic resistance relationships. Enhanced carbohydrate metabolism, nucleotide biosynthesis, and particularly putrescine degradation pathways indicate a coordinated metabolic strategy that prioritizes energy efficiency and cellular protection (Schneider et al., 2013). The strategic optimization of energy metabolism through preserved ATP synthase and NADH dehydrogenase complex function demonstrates that evolved bacteria develop superior mechanisms for maintaining electron transport chain efficiency while minimizing ROS generation—a critical adaptation given that rising temperatures naturally increase oxidative stress burden (Sies & Jones, 2020). This metabolic efficiency is further evidenced by the reduced reliance on traditional ROS scavenging enzymes, suggesting that evolved strains achieve oxidative stress tolerance through preventive mechanisms rather than reactive detoxification, thereby conserving metabolic resources for growth and adaptation under multiple stressors (Ren et al., 2020).

### 4.2. Cross-Protection Mechanisms and Environmental Management Implications

The most significant finding of this study is the demonstration that environmental stress adaptations confer cross-protection against antimicrobial challenges through shared regulatory pathways and metabolic networks. The observed reduction in antibiotic resistance gene expression coupled with enhanced tolerance in evolved strains challenges traditional assumptions about resistance mechanisms and reveals a more nuanced relationship between environmental stress and antimicrobial resistance (Chen et al., 2019). The downregulation of broad-spectrum resistance regulators like marA alongside maintained cellular integrity through improved cell wall synthesis components suggests that bacteria can achieve antibiotic tolerance through structural and metabolic optimization rather than energy-intensive resistance gene overexpression. This finding is particularly relevant given that antibiotic treatment can induce ROS levels reaching 150-200% above baseline, with complex cellular responses that differ significantly from traditional oxidative stress responses (Ye et al., 2024; Dwyer et al., 2014).

Our proteomic validation experiments provide critical evidence that oxygen availability serves as a master regulator coordinating cellular systems to optimize survival under aerobic stress while reducing the metabolic burden of maintaining high-level antimicrobial resistance. The comprehensive reconfiguration of resistance mechanisms under elevated oxygen conditions—including downregulation of penicillin-binding proteins, modification of membrane architecture, and recalibration of two-component regulatory systems—demonstrates that environmental management strategies targeting oxygen levels could effectively mitigate antibiotic resistance proliferation in aquatic ecosystems (García et al., 2020). This oxygen-dependent regulation aligns with previous observations that bacteria must balance energy conservation with defense capabilities, often shifting from constitutive to inducible resistance mechanisms when resources become limited (Kalia et al., 2020). The ability of evolved strains to maintain antibiotic tolerance with reduced genetic investment represents an evolutionary fine-tuning that could be exploited for environmental management, where controlled oxygen perturbations might disrupt the energy-intensive pathways that support antibiotic resistance while preserving essential cellular functions (Zhu et al., 2023).

These discoveries have profound implications for addressing the climate warming-antibiotic resistance nexus in natural aquatic systems. The identification of oxygen modulation as a potential strategy for mitigating temperature-induced ARG expression provides a nature-based solution that could complement traditional disinfection approaches, which often face limitations including high costs, energy intensity, and secondary environmental impacts (Department et al., 2020). The regulatory networks identified in this study—particularly the oxyR-metR and arcA-betI systems—represent potential targets for developing interventions that could disrupt bacterial stress adaptation mechanisms without relying on chemical treatments that may further select for resistant strains (Alaoui Mdarhri et al., 2022). Future research should investigate whether these coordinated stress response mechanisms are conserved across diverse bacterial species in natural environments and explore the long-term evolutionary stability of these adaptations under sustained environmental pressure. Additionally, translating these mechanistic insights into practical oxygen management strategies for controlling antibiotic resistance in warming aquatic ecosystems represents a critical next step for addressing this global health challenge in an era of accelerating climate change (Li et al., 2022).

**Figure 7.**
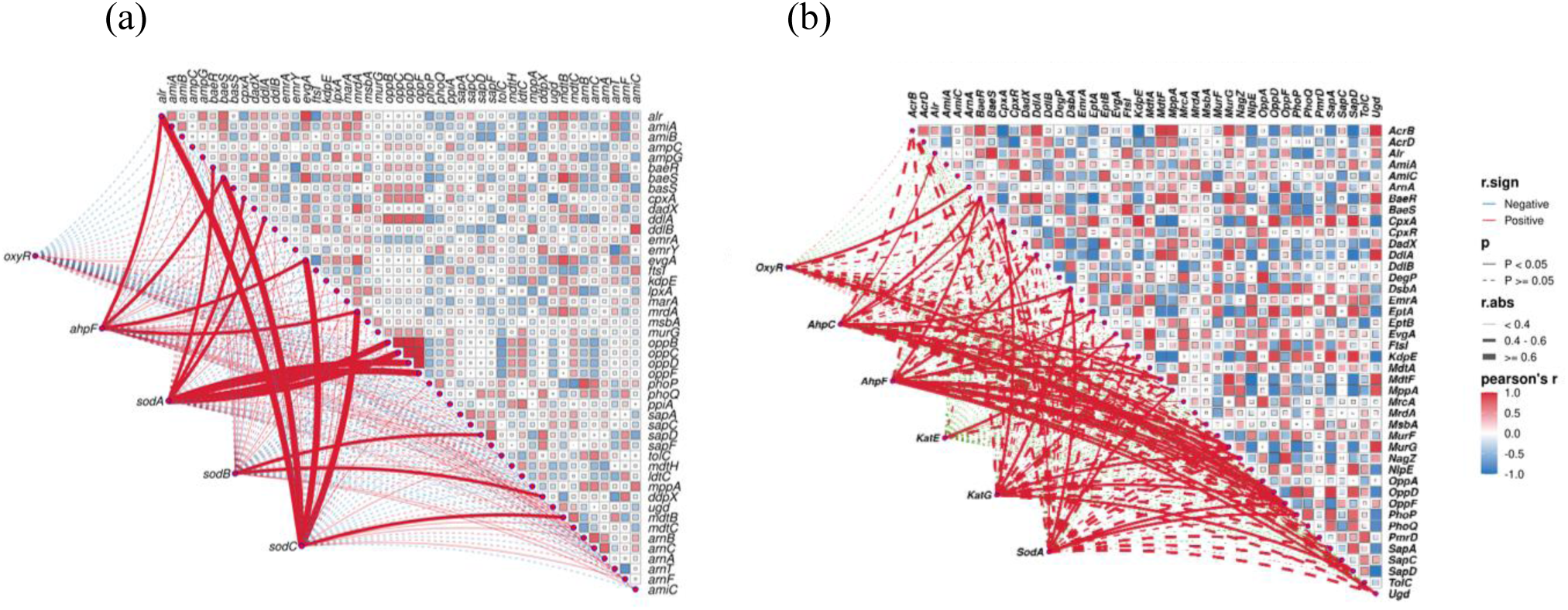
A network analysis depicting the correlations between ROS scavengers and ARGs in *E. coli* based on (a) transcriptomics data and (b) proteomics data.

## 5. Conclusion

This study elucidates sophisticated regulatory and metabolic adaptations enabling *E. coli* to survive high temperature stress through integrated transcriptional rewiring. Evolved strains demonstrate a fundamental shift from reactive damage repair to proactive resilience through three adaptations: optimized energy metabolism via preserved ATP synthase and NADH dehydrogenase function, strategic ROS management prioritizing metabolic efficiency, and streamlined antibiotic resistance maintaining tolerance while reducing metabolic burden. The ability to maintain antibiotic tolerance with reduced ARG expression demonstrates bacterial potential for efficient survival under dual climate-antimicrobial pressures. These discoveries identify oxygen availability as a potential modulator of temperature-induced ARG expression, providing novel targets for environmental management strategies mitigating antibiotic resistance proliferation in warming aquatic ecosystems.

## Acknowledgment

This work was supported by the Marsden Fund Council, administered by the Royal Society Te Apārangi, New Zealand [Grant No. MFP-UOA2018].

## May use

Acquiredresistance mechanisms mediated by the bacterial cell wall, nucleic acids, and proteins play a pivotal role in the genesis of MDR. Bacteria can modify their cell wall structure, produce resistant enzymes, exhibit mutations in antibiotic-targetedgenes, and acquire resistant genes through horizontal gene transfer.

(A Comprehensive Review of Molecular Mechanisms Leadingto the Emergence of Multidrug Resistance in Bacteria)

The coordinated upregulation of these metabolic pathways in our evolved strains suggested the development of an integrated stress response network that builds upon the foundational regulatory mechanisms identified in our regulon analysis. This metabolic reprogramming, particularly the enhancement of polyamine catabolism, appears to be a key evolutionary adaptation that enables evolved strains to better manage multiple environmental stressors through enhanced versions of the regulatory pathways we characterized.

